# Accurate *de novo* prediction of RNA 3D structure with transformer network

**DOI:** 10.1101/2022.10.24.513506

**Authors:** Chenjie Feng, Wenkai Wang, Renmin Han, Ziyi Wang, Lisa Ye, Zongyang Du, Hong Wei, Fa Zhang, Zhenling Peng, Jianyi Yang

## Abstract

RNA 3D structure prediction remains challenging though after years of efforts. Inspired by the recent breakthrough in protein structure prediction, we developed trRosettaRNA, a novel deep learning-based approach to *de novo* prediction of RNA 3D structure. Like trRosetta, the trRosettaRNA pipeline comprises two major steps: 1D and 2D geometries prediction by a transformer network; and full-atom 3D structure folding by energy minimization with constraints from the predicted geometries. We benchmarked trRosettaRNA on two independent datasets. The results show that trRosettaRNA outperforms other conventional methods by a large margin. For example, on 25 targets from the RNA-Puzzles experiments, the mean RMSD of the models predicted by trRosettaRNA is 5.5 Å, compared with 10.5 Å from the state-of-the-art human group (i.e., Das). Further comparisons with two recently released deep learning-based methods (i.e., DeepFoldRNA and RoseTTAFoldNA) show that all three methods have similar accuracy. However, trRosettaRNA yields more accurate and physically more realistic side-chain atoms than DeepFoldRNA and RoseTTAFoldNA. Finally, we apply trRosettaRNA to predict the structures for the Rfam families that do not have known structures. Analysis shows that for 263 families, the predicted structure models are estimated to be accurate with RMSD < 4 Å. The trRosettaRNA server and the package are available at: https://yanglab.nankai.edu.cn/trRosettaRNA/.

## INTRODUCTION

Similar to proteins, RNA molecules’ biological function is determined by their 3D structures. However, most RNA structures are highly flexible ^1^, making them difficult to be solved by experiments. For example, only ~6000 RNA structures are deposited in the Protein Data Bank (PDB) ^2^, which is much less than the number of deposited protein structures (~190000). Thus, there is a great demand for developing efficient algorithms to predict RNA 3D structures.

The current RNA 3D structure prediction methods can be divided into two groups: template-based methods and *de novo* methods. Template-based methods predict the target structure using homologous templates in PDB. For example, the representative methods, such as ModeRNA ^3^ and MMB ^4^, work by reducing the structure searching space with homologous structures. In general, the predicted structure models by template-based methods are accurate when homologous templates exist in PDB. However, the progress for template-based methods is slow, due to the limited number of known RNA structures and the difficulty of aligning RNA sequences.

On the contrary, *de novo* methods attempt to construct the lowest-energy 3D conformations by simulating the folding process from scratch. With molecular dynamic simulations and/or fragment assembly, methods such as FARFAR ^5^, FARFAR2 ^6^, SimRNA ^7^, and 3dRNA ^8, 9^, work well for small RNAs (<100 nucleotides). Nevertheless, it is hard to generate accurate 3D structures for large RNAs with complicated topologies with such methods, due to the inaccurate force field parameters and the huge space of 3D conformations. To partly address this issue, inter-nucleotide contacts predicted by direct coupling analysis (DCA) have been used to guide the structure simulations to improve the modeling accuracy ^10–12^. In addition, given the hierarchical nature of RNA structure folding, RNAComposer ^13^, Vfold ^14^, and MC-Fold ^15^ derive 3D structures from secondary structures. They are very fast but the modeling accuracy largely depends on the quality of the input secondary structures. The RNA-Puzzles experiments indicate that it remains a grand challenge to accurately predict the structures for large RNAs with complex architectures ^16, 17^.

Deep learning has recently been used to improve *de novo* RNA 3D structure prediction. The predicted inter-nucleotide contacts by the residual convolutional network (ResNet) are about two times more accurate than DCA, improving 3D structure prediction to some extent ^18, 19^. It was shown that with the model selection from a geometric deep learning-based scoring system (ARES), the FARFAR2 protocol predicted the most accurate models for four targets in the blind test of the RNA-Puzzles experiments ^20^. Recently, significant progress was reported in DeepFoldRNA ^21^ and RoseTTAFoldNA ^22^, which are mostly inspired by the success of AlphaFold2 in protein structure prediction ^23^.

In this work, we introduce trRosettaRNA, a novel deep learning-based *de novo* approach to RNA 3D structure prediction. It is partly inspired by the successful application of deep learning in protein structure prediction, especially in AlphaFold2 ^23^ and our previous method trRosetta ^24–26^. Benchmark tests indicate that trRosettaRNA significantly advances the accuracy of RNA 3D structure prediction over traditional methods. Compared to DeepFoldRNA and RoseTTAFoldNA, trRosettaRNA yields more accurate and physically more realistic side-chain atoms. Finally, we use trRosettaRNA to predict the structures for the Rfam families with unknown structures.

## RESULTS

### Overview of trRosettaRNA

The architecture of trRosettaRNA is depicted in Figure 1A. Starting from the nucleotide sequence of an RNA of interest, a multiple sequence alignment (MSA) and a secondary structure are first generated by the programs rMSA (https://github.com/pylelab/rMSA) and SPOT-RNA ^27^, respectively. They are then converted into an *MSA representation* and a *pair representation*, which are fed into a transformer network (named RNAformer, see Figure 1B and Methods for more details) to predict 1D and 2D geometries (see Figure S1). Similar to trRosetta, these geometries are converted into restraints to guide the last step of 3D structure folding based on energy minimization (see Methods).

**Figure 1.**
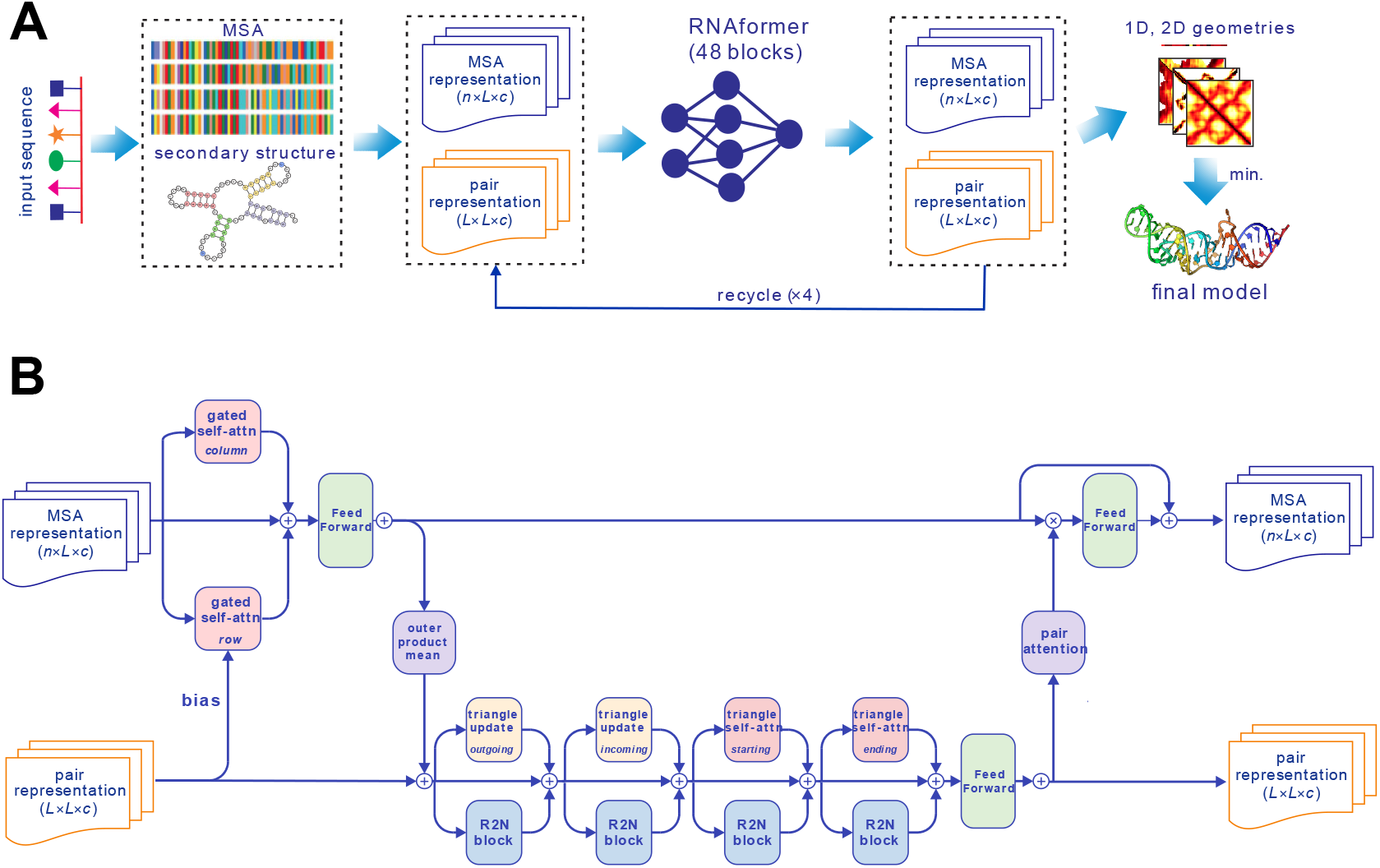
Overall architecture of trRosettaRNA. (A) flowchart of trRosettaRNA. (B) structure of each RNAformer block. *n, L*, and *c* are the number of sequences in the MSA, the length of the query sequence, and the number of channels, respectively.

### Performance of trRosettaRNA on 101 independent RNAs

To evaluate trRosettaRNA, we collected 101 non-redundant RNA structures that are released after the date of the training RNAs (i.e., 2017-01). trRosettaRNA achieves an average RMSD of 4.0 Å on these RNAs, for which 74.2% are predicted with RMSD < 4 Å.

We compare trRosettaRNA with two representative *de novo* methods, RNAComposer ^13^ and SimRNA ^7^. The same secondary structures (from SPOT-RNA) were fed into both methods for fair comparisons. For RNAComposer, we submitted the RNA sequences and the secondary structures to its web server to collect the predictions. SimRNA was installed and run locally in our computer cluster. Only 93 RNAs are used in this comparison because RNAComposer failed to return results for eight RNAs. On these 93 RNAs, the average RMSD by trRosettaRNA (4.0 Å) is significantly lower than those by RNAComposer (18.8 Å; P-value = 2.1E-37) and SimRNA (19.2 Å; P-value = 2.7E-38). trRosettaRNA outperforms RNAComposer and SimRNA for 92 and 93 cases, respectively (Figure 2A). 73.1% of the models predicted by trRosettaRNA are with RMSD < 4 Å, significantly higher than RNAComposer (1.1%) and SimRNA (0%). These data demonstrate the superiority of the proposed pipeline over traditional RNA structure prediction methods.

**Figure 2.**
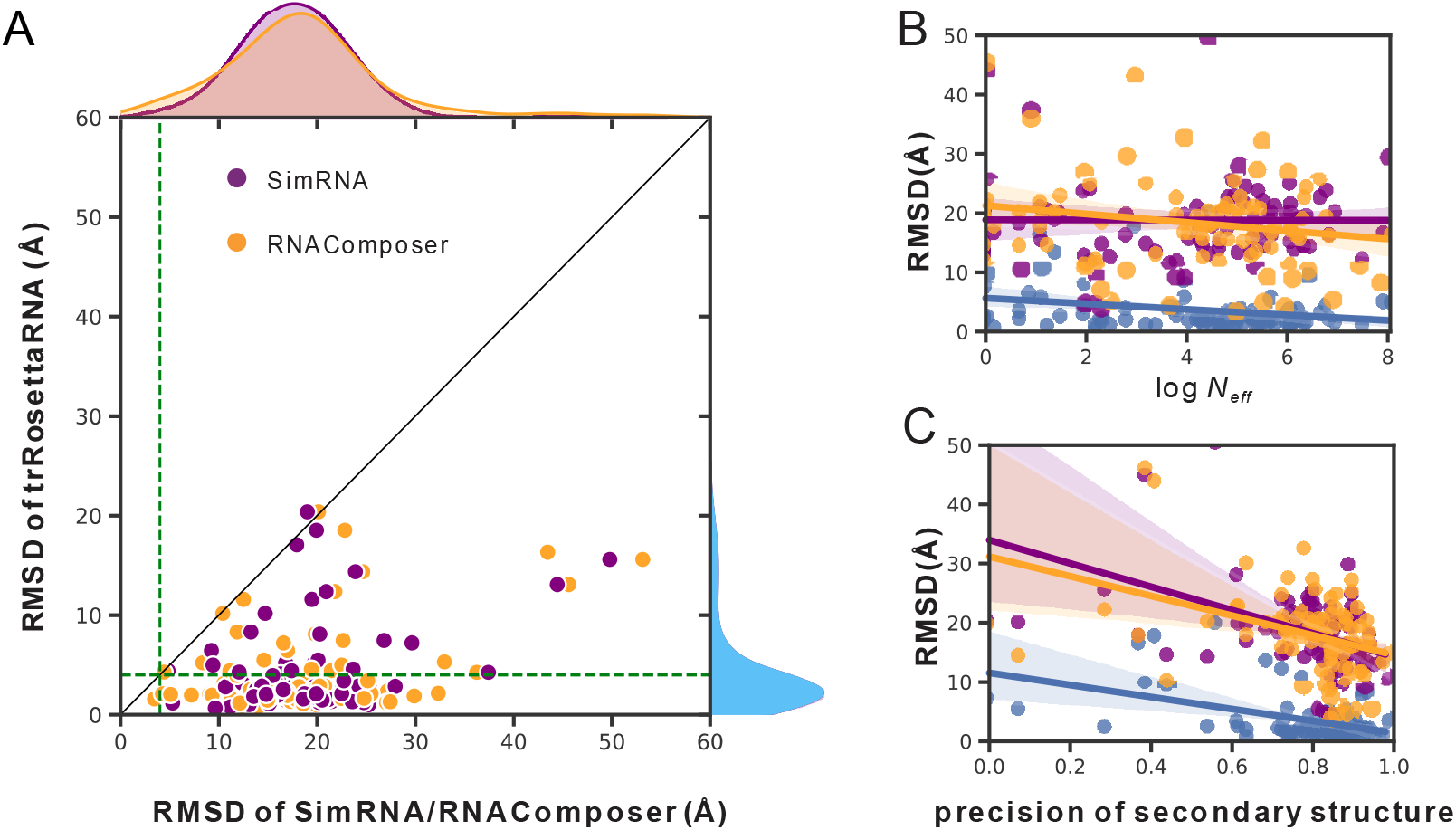
Performance on 93 RNAs. (A) head-to-head comparison between trRosettaRNA and two representative methods, SimRNA and RNAComposer. The dashed horizontal and vertical lines correspond to an RMSD of 4 Å. The bar plots show the RMSD distributions. (B) the RMSD as a function of the logarithm of the MSA depth (*N*_*eff*_). (C) RMSD as a function of the precision of the predicted secondary structure. The blue, purple, and orange dots in B and C refer to trRosettaRNA, SimRNA, and RNAComposer, respectively.

To investigate why trRosettaRNA could generate accurate structure models, we analyze the impact of the input features (i.e., MSA and secondary structure). The MSA quality is measured by the depth, i.e., the logarithm of the effective number (*N*_*eff*_) of homologous sequences at 80% sequence identity. As shown in Figure 2B, the RMSD of the trRosettaRNA model is slightly correlated with *N*_*eff*_ (Pearson correlation coefficient, PCC = −0.29). trRosettaRNA outperforms RNAComposer and SimRNA at all *N*_*eff*_ levels, especially on targets with high *N*_*eff*_ values. For 82.1% of the 55 RNAs with *N*_*eff*_ >50, trRosettaRNA can predict accurate structure models with RMSD < 4 Å. As expected, the RMSDs of the models by RNAComposer and SimRNA have a weak correlation with the MSA depth (PCCs are −0.19 and −0.003, respectively), probably because they do not use MSA during modeling. In contrast, there is a strong correlation between the RMSDs of the predicted models and the precision of the predicted secondary structures for all methods (PCCs are −0.389, −0.427, and −0.287 for trRosettaRNA, SimRNA, and RNAComposer, respectively, Figure 2C). This is consistent with the observations that precise RNA secondary structure prediction plays a key role in successful 3D structure modeling ^16, 17^.

Note that the above test RNAs are selected based on the release date but their sequence similarity to the training RNAs is not considered. We check whether the high accuracy is due to their sequence redundancy with the training set. For 60 out of the 101 RNAs, no similar sequences can be detected in the training set according to the BLASTN ^28^ search at an E-value cutoff of 10. The mean RMSD on these RNAs is 4.4 Å, compared to 3.5 Å on the remaining 41 RNAs that have similar RNAs in the training set. The difference is however not significant statistically (P-value = 0.27; Figure S2). We thus conclude that the results for these 101 RNAs are not over-estimated.

Figure 3 shows the results for three example RNAs from the above dataset. High-quality MSAs (*N*_*eff*_ >50) were generated for these RNAs. SPOT-RNA predicted accurate secondary structures (precision > 0.9, Figures 3A, B) for two RNAs but a poor secondary structure for the remaining one (precision = 0.44, Figure 3C). The RMSDs of the models predicted by SimRNA and RNAComposer are higher than 10 Å for these three RNAs. In contrast, the trRosettaRNA models are very accurate for the first two RNAs (RMSD < 2Å). For the last RNA with poor secondary structure prediction, the predicted model by trRosettaRNA is still satisfactory with 2.9 Å RMSD. This shows that the incorporation of MSA, secondary structure, and transformer network can significantly advance the accuracy of RNA 3D structure modeling.

**Figure 3.**
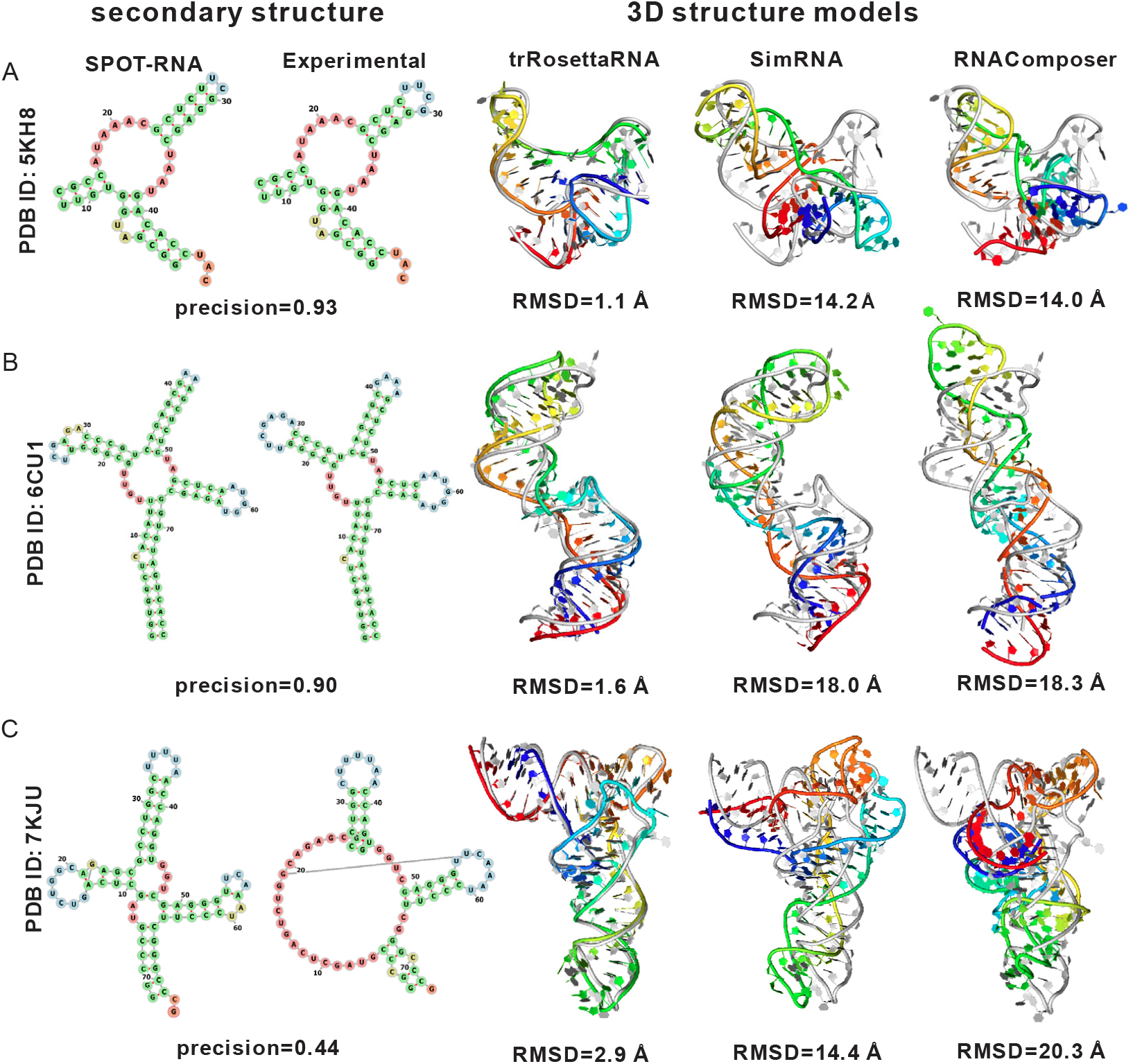
Case study on three example targets. Left: comparison between the secondary structures predicted by SPOT-RNA and those extracted from experimental structures. Right: comparison between the 3D structures predicted by trRosettaRNA, SimRNA, and RNAComposer. The predicted structures (rainbow cartoons) are superposed to the experimental structures (gray cartoons).

### Performance of trRosettaRNA on RNA-Puzzles targets

We further test trRosettaRNA on 30 targets from the RNA-Puzzles experiments ^16, 17^. The target information and the prediction results are summarized in Table S1. These targets are harder to predict than the 101 RNAs, as revealed by the increased value of RMSD (5.7 Å). Nevertheless, more than half of these targets (18/30, shown in Figure S3 and highlighted in bold in Table S1) are predicted with < 4 Å RMSD, indicating the effectiveness of our method.

We compare the trRosettaRNA predictions with the original submissions from the RNA-Puzzles experiments. According to the official assessments ^16, 17^, the state-of-the-art approach is the Das group, which submitted models for 25 targets. Table S1 summarizes the results on these targets for the first model from the Das group (denoted by Das_1) and the best of the first models from all groups (denoted by Best_1). The average RMSD of the models by our method is 5.5 Å, compared with 10.5 Å and 8.4 Å from Das_1 and Best_1, respectively. Note that 16 of the 25 models (64%) by trRosettaRNA have < 4 Å RMSDs, compared with 1 and 6 for Das_1 and Best_1, respectively. These data suggest the superiority of trRosettaRNA over other methods. Nevertheless, there are also a few targets for which trRosettaRNA models are less accurate than the Best_1 models. These are special RNA targets, such as in free conformations (e.g., PZ14Free) or have complicated interactions with protein (e.g., PZ30). Note that some groups utilized human experts and/or partial experimental data to guide the modeling during the prediction seasons of the RNA-Puzzles experiments. In contrast, the trRosettaRNA predictions are fully automated.

### Comparison with DeepFoldRNA

During the preparation of this manuscript, another approach (named DeepFoldRNA) with a similar idea was proposed ^21^. We compare trRosettaRNA with DeepFoldRNA on the above datasets. DeepFoldRNA was installed and run locally with default options (using the same secondary structure and MSA). On the 30 RNA-Puzzle targets (Figure 4A), the average RMSD of the DeepFoldRNA models is 5.1 Å compared to 5.7 Å for our method. The RMSD difference between both methods is not significant (P-value = 0.59). We further check the ratio of accurate models with RMSD < 4 Å. At this cutoff, 60% of our predictions are accurate, slightly higher than that of DeepFoldRNA (50%). On the other dataset of 101 RNAs (Figure 4B), the average RMSD of trRosettaRNA is 4.0 Å, 0.2 Å lower than the DeepFoldRNA (4.2 Å; P-value = 0.82). Note that both methods are inspired by the recent advance in protein structure prediction with a similar idea, which explains the close performance.

**Figure 4.**
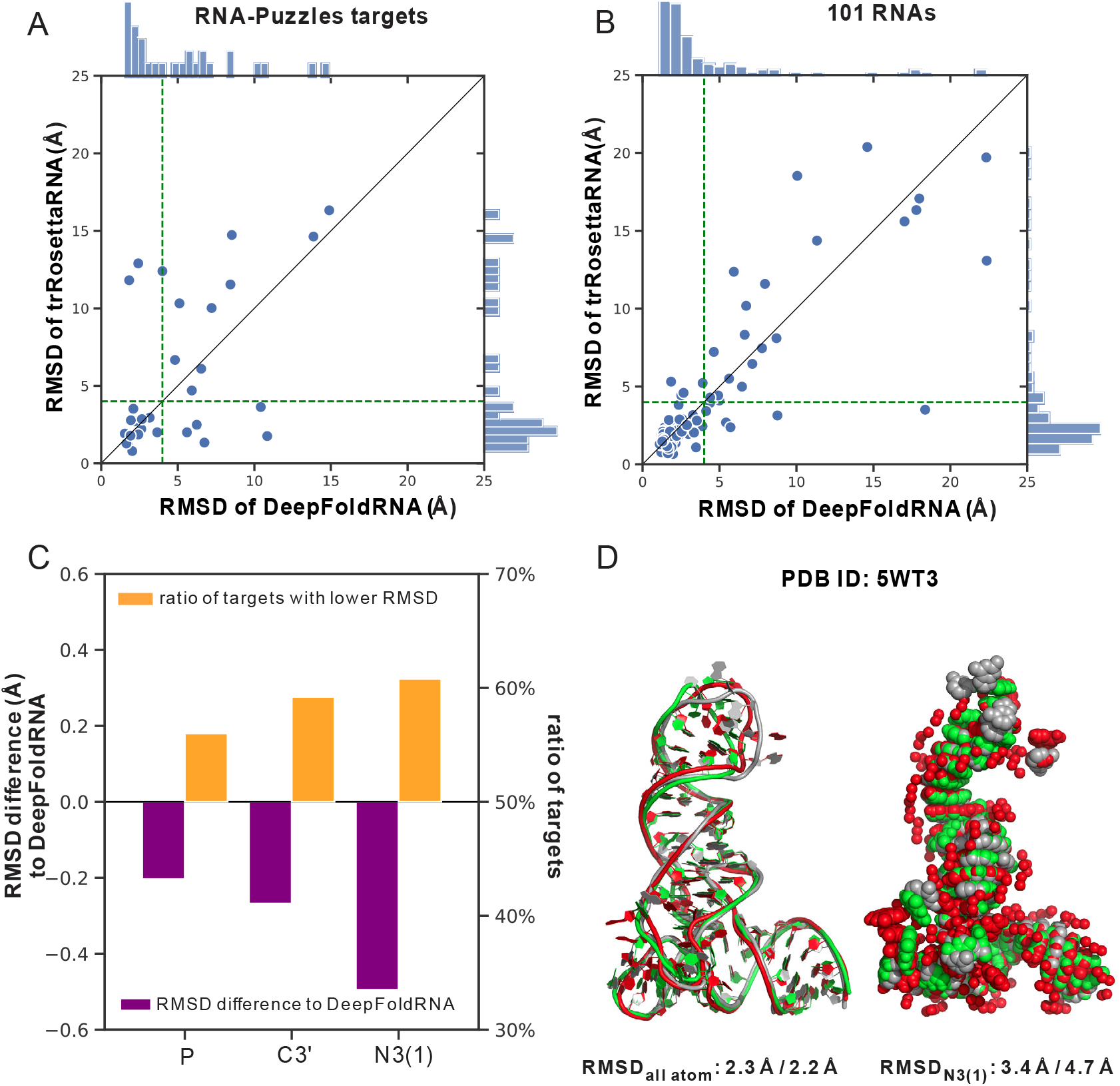
Comparison between trRosettaRNA and DeepFoldRNA. (AB) head-to-head RMSD comparison between trRosettaRNA and DeepFoldRNA on (A) 30 RNA-Puzzles targets and (B) 101 non-redundant RNAs. The dashed horizontal and vertical lines correspond to an RMSD of 4 Å. The bar plots show the RMSD distributions. (C) comparison of RMSDs calculated based on different atoms on all test targets. The purple bars refer to the average RMSD difference between the two methods (the lower this value is, the more significantly trRosettaRNA outperforms DeepFoldRNA). The orange bars refer to the ratio of targets on which trRosettaRNA outperforms DeepFoldRNA. (D) an example RNA (PDB ID: 5WT3) for which trRosettaRNA builds more precise side chains than DeepFoldRNA. Left: the superposition of predicted structures (green for trRosettaRNA, red for DeepFoldRNA) to the experimental structures (gray). Right: the same with left except that only the nucleobases are shown. The RMSD comparisons are shown in the “trRosettaRNA/DeepFoldRNA” format.

The main methodological difference between both methods is in the construction of the atomic structure. In DeepFoldRNA, coarse-grained models are first built from predicted restraints. The remaining atoms are then added and refined by SimRNA and QRNA ^29^. In contrast, trRosettaRNA directly optimizes full-atom models with 1D and 2D restraints from the network RNAformer. As shown in Figure 4C, both methods perform on par when the backbone atom (P) is assessed only. However, trRosettaRNA shows superiority when other side-chain atoms are considered. For example, for the N3(1) atom (i.e., N3 for C, U; N1 for A, G), trRosettaRNA outperforms DeepFoldRNA for 60.8% of targets. Figure 4D shows a representative target (PDB ID: 5WT3) for which both methods generate high-quality 3D structures with full-atom RMSD < 2.5 Å. However, the side-chain atoms of the trRosettaRNA model are closer to the experimental structure than the DeepFoldRNA model (RMSD_N3(1)_ 3.4 Å vs 4.7 Å).

To summarize, trRosettaRNA performs similarly to DeepFoldRNA when measured by all-atom RMSD; but outperforms DeepFoldRNA when the side-chain atoms are measured.

### Comparison with RoseTTAFoldNA

While submitting this manuscript, another deep learning-based method (named RoseTTAFoldNA) was released ^22^. We further compare our method with RoseTTAFoldNA. RoseTTAFoldNA can predict the structures for both nucleic acids and protein-nucleic acids complexes. As its training set consists of all the PDB structures released before 2020-05-01, 38 out of the 101 RNAs released after this date are used for comparison. The RNA-Puzzles targets are also not compared here due to the same reason. RoseTTAFoldNA was installed and run locally with default options (using the same MSA). On these 38 RNAs, the average RMSD of RoseTTAFoldNA (5.0 Å) is higher than trRosettaRNA (3.8 Å). The RMSD difference between both methods is not significant (P-value = 0.36). The higher average RMSD of RoseTTAFoldNA is mainly due to its poor performance on two RNAs (40.7 Å on 7LVA, 21.2 Å on 6LVR, highlighted in Figure S4). As shown in Figure S5, a few key interactions are missed in the RoseTTAFoldNA models for both RNAs. In addition, there are some structural violations (e.g., the chain break highlighted by the circle in Figure S5A) in the RoseTTAFoldNA model for 7LVA, which may be because its end-to-end prediction is not refined. Nevertheless, for the remaining RNAs, RoseTTAFoldNA has a similar average RMSD (3.6 Å) to trRosettaRNA (3.4 Å, P-value=0.74).

### Improved prediction of inter-nucleotide contacts

Accurate predictions of inter-nucleotide contacts are helpful for successful 3D structure prediction ^10–12^. We further compare trRosettaRNA with two deep learning-based inter-nucleotide contact prediction methods, namely, RNAcontact ^18^ and SPOT-RNA-2D ^19^. A total of 26 and 12 RNAs are used to compare with RNAcontact and SPOT-RNA-2D, respectively, by the consideration of the similarity to their training RNAs. As shown in Figure S6, the long-range contacts predicted by trRosettaRNA are more accurate than both two methods on almost all compared targets (83.3% for RNAcontact, 92.3% for SPOT-RNA-2D). Note that the network ResNet ^30^ was applied in both methods. The above data demonstrate the advantage of the transformer network (RNAformer). We use PZ20 (PDB ID: 5Y85), a ribozyme with a four-way junction fold, to investigate how RNAformer works. For this target, we extract the row-wise attention maps from the last block of RNAformer. As shown in Figure S7A, some attention maps (highlighted by red frames) show similar patterns to the true contact map, providing rich interaction signals beyond the base-pairing information in the secondary structure. It turns out that RNAformer detects more inter-nucleotide contacts than the other two ResNet-based methods (Figure S7B). As a result, the 3D structure model predicted by trRosettaRNA is very accurate with 2.0 Å RMSD (Figure S7C).

### Impact of the predicted 1D and 2D geometries

The geometries predicted by the RNAformer network consist of 1D orientations and 2D contacts, distances, and orientations (Figure S1). To analyze their contributions, we compare the modeling results using different geometries on the 30 RNA-Puzzles targets (Figure 5A and Table S2). Using the 2D distance restraints only, trRosettaRNA achieves a reasonable RMSD of 6.1 Å. This value is reduced to 5.8 Å when the 1D and 2D orientations are included. Furthermore, with the help of 2D contacts, the RMSD drops to 5.7 Å. We use the target PZ19 (PDB ID: 6TB7) to show the impacts of different restraints. As shown in Figure 5B, using 2D distance restraints only, trRosettaRNA can generate a reasonable structure model (RMSD < 4 Å). But the loop region (highlighted by the black square in Figure 5B) is wrongly twisted and far from the helix. The introduction of 1D and 2D orientations slightly fixes the wrong twist of this region. The 2D contact restraints force the loop nucleotides to move closer to the helix, resulting in a more accurate model with 1.6 Å RMSD.

**Figure 5.**
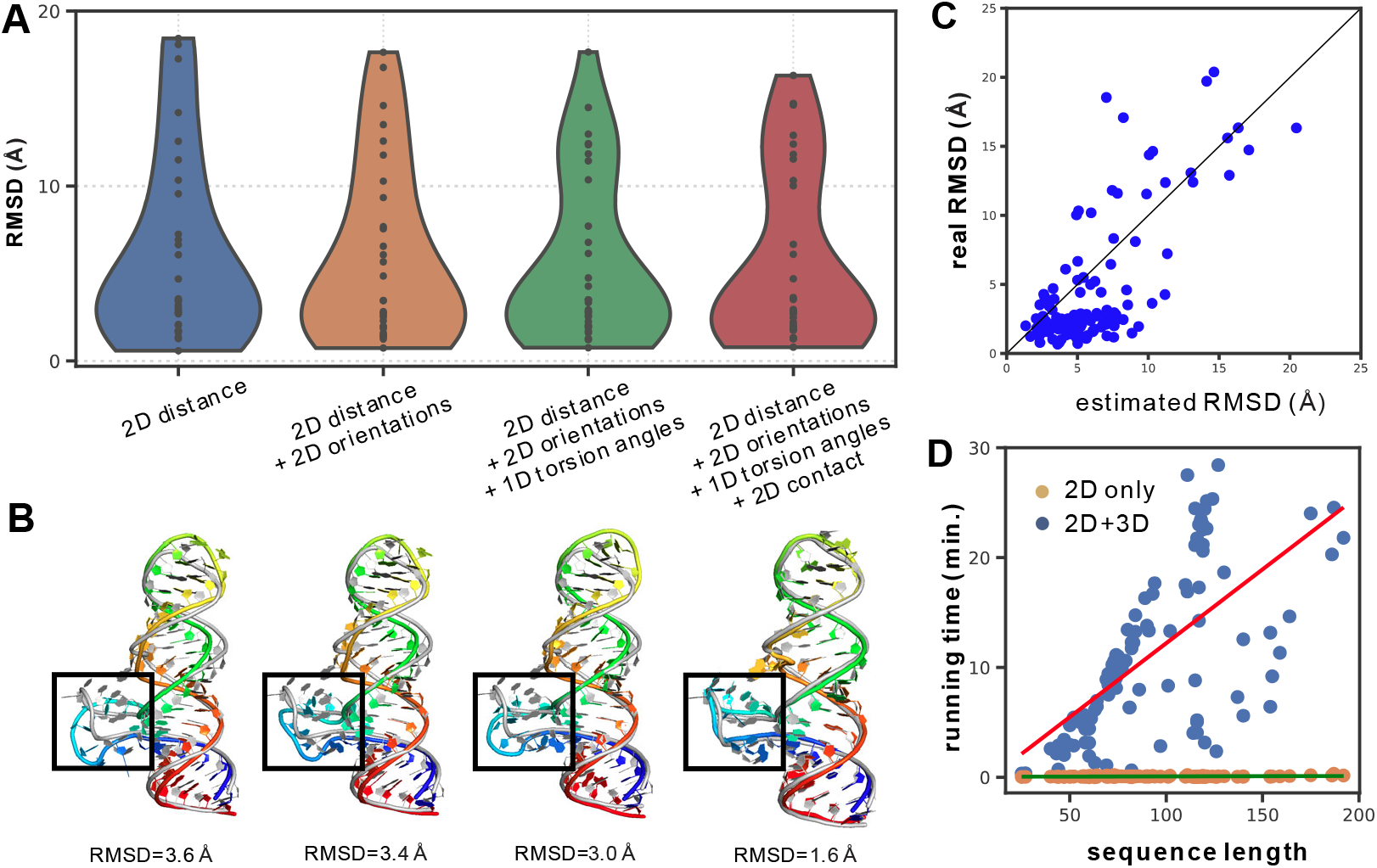
Summary of the folding results by different restraints. (A) contribution of the various restraints to the trRosettaRNA modeling accuracy in terms of the RMSD for the 30 RNA-Puzzles targets. The *x*-axis of the violin plot signifies the different restraints. (B) an example (PZ29) to illustrate the impact of different restraints. The predicted models (rainbow cartoon) are superposed to the experimental structures (gray cartoon). (C) head-to-head comparison between the estimated and real RMSD for all RNAs in the benchmark datasets. (D) the relationship between the running time and the sequence length. The 2D geometry predictions (orange dots) were run on 1 GPU card. The 3D folding was performed on 1 CPU core.

### Confidence score of the predicted structure models

To guide real-world application, the confidence scores of the predicted protein structure models have been estimated reliably in trRosetta ^24–26^. A similar estimation can be extended to trRosettaRNA. Specifically, we first calculate a few variables reflecting the confidence of the predicted distance maps and the convergence of the top structure models (see Methods for more details). Then a linear regression on these variables is employed to fit the RMSD values. For the RNAs from the benchmark datasets, the estimated RMSDs (eRMSDs) correlate well with the real RMSDs of the predicted models (PCC = 0.76, Figure 5C).

### Analysis of the running time

We decompose the running time of trRosettaRNA into two parts: geometry prediction and 3D structure generation. The time for MSA generation is not discussed here as it can be flexible depending on the searching algorithms and sequence databases. Figure 5D shows that trRosettaRNA spends most time in the generation of 3D structure (> 95%). With the increase in sequence length, the running time increase linearly. In general, it takes < 30 minutes to complete the prediction for a typical RNA with < 200 nucleotides.

### Application to Rfam families with unknown structures

It remains challenging to solve RNA structures by experiment. For example, only 123 out of the 3938 families in the Rfam database (version 14.4) have experimentally resolved 3D structures ^31^. We sought to predict the structures for the Rfam families that have no experimental structures. We collected 1752 unsolved families that are 50-200 nucleotides long and have more than 30 members. For each family, we use its consensus secondary structure along with the MSA derived from the consensus sequence as the input features to trRosettaRNA. Most of these families are not predicted well, with eRMSD > 10 Å for 975 out of 1752 families. This may reflect the difficulty of determining the structures for these families.

Nevertheless, trRosettaRNA does predict accurate structures for 263 families with eRMSD < 4 Å. For 27 of these families, the predicted structure models are of novel topologies, i.e., they do not have any similar structures in PDB according to the program RNAalign (TM-score_RNA_^32^ ≥ 0.45). In Figure 6B, we show the predicted structures for 12 families with distinct topologies, among which the last six are of novel topologies. These high-confidence models are anticipated to provide a structural basis for understanding their biological functions and guide their experimental determinations. For example, for the family hammerhead ribozyme (RF02275), which catalyzes reversible cleavage and ligation reactions at a specific site within an RNA molecule, the trRosettaRNA model is estimated to have 0.6 Å RMSD. The trRosettaRNA models for the 263 families with eRMSD < 4 Å are available on our website.

**Figure 6.**
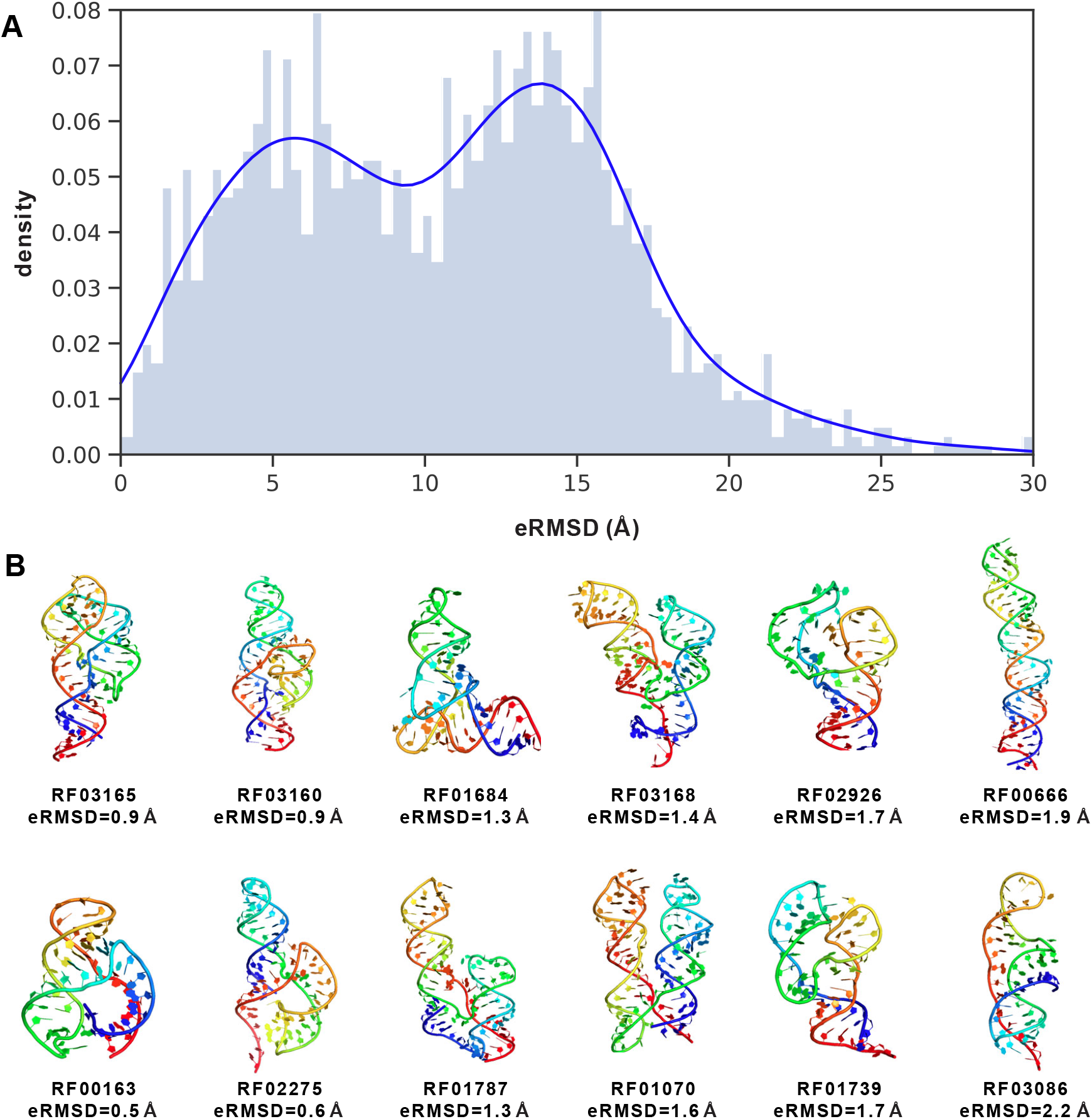
Application of trRosettaRNA to Rfam families with unknown structures. (A) eRMSD distributions of the predicted structure models. (B) 12 selected example families with eRMSD < 4Å.

## CONCLUSIONS

We have developed trRosettaRNA, a *de novo* approach to the accurate prediction of RNA 3D structure with the transformer network. trRosettaRNA follows trRosetta’s two-step procedure, one is for 1D and 2D geometries prediction, and the other is for 3D structure generation by energy minimization. With the transformer network in the first step, the 1D and 2D geometries are predicted with high accuracy, which are translated into accurate 3D structure models in the second step. We have comprehensively benchmarked trRosettaRNA on two independent datasets. It shows that trRosettaRNA predicts significantly more accurate models than the conventional methods. In addition, trRosettaRNA performs similarly to the recent deep learning-based methods DeepFoldRNA and RoseTTAFoldNA in terms of all-atom RMSD; but yields more accurate and physically more realistic side-chain atoms. Finally, trRosettaRNA is applied to predict the structures for Rfam families with no experimental structures. For 263 families, the predicted structure models are estimated to be accurate with < 4 Å RMSD. Note that these only represent a small proportion of the current Rfam families. We believe that more families would be predicted with high accuracy in the future, by the consistent improvement of algorithms and the deposition of more experimental RNA structures.

## METHODS

### 1. trRosettaRNA algorithm

As shown in Figure 1A, the full pipeline of trRosettaRNA consists of three major steps: preparation of input data, prediction of 1D and 2D geometries, and generation of 3D structure.

#### Step 1. Preparation of input data

For a given query RNA, the first step of trRosettaRNA is to prepare an MSA and a secondary structure. Two different MSAs are generated for each query sequence. The first is generated by using the program rMSA against multiple sequence databases (NCBI’s nt, Rfam, and RNAcentral ^33^). The second is obtained by running the program Infernal ^34^ against the smaller database RNAcentral with two iterations, which is very fast. Then we select the final MSA based on the qualities of the predicted distance maps (measured by the average of standard deviations of the probability values of each nucleotide pair, Figure S8). The secondary structure is predicted by SPOT-RNA ^27^ from the query sequence. Here we use the predicted probability matrix as the input, which contains more information than the dot-bracket representation.

#### Step 2. Prediction of 1D and 2D geometries

The second step of trRosettaRNA is to predict the 1D and 2D geometries by deep learning. We design a transformer network (named RNAformer) similar to the network Evoformer in AlphaFold2. At the very start, the input MSA and secondary structure are converted into two representations, i.e., the MSA representation (i.e., MSA embedded by nucleotide types) and the pair representation (including the direct couplings derived from MSA and the probability matrix of the predicted secondary structure). We adopt a transformer-based module (i.e., RNAformer) to update both representations. More specifically, each block of RNAformer can be divided into four steps according to the update direction (Figure 1B).

**1) MSA to MSA**. To update the MSA representation by itself, we perform row- and column-wise gated self-attention operations and combine the corresponding results. A feed-forward layer is employed to introduce nonlinearity. Note that the pair information participates in the row-wise attention by adding bias to the attention maps.

**2) MSA to pair**. We perform an outer product operation on the self-updated MSA representation to transform it into the pair format. In detail, the MSA representation is linearly projected to a smaller dimension. Then for the nucleotide pair (*i, j*), the outer products of the vectors from the *i*-th and the *j-*th columns of the MSA representation are averaged over the homologous sequences to update the representation for this pair.

**3) Pair to pair**. After the above step, we perform the triangle updates, followed by a feed-forward layer. For each triangle update layer, we use a multi-scale network Res2Net ^35^ to enhance the ability to model the local details.

**4) Pair to MSA**. The updated pair representation is then linearly projected to the pair-wise attention maps, which are then multiplied on the MSA representation, followed by a feed-forward layer.

A single-pass RNAformer consists of 48 blocks, which are cycled 4 times in the complete inference (Figure 1A). The final predicted probability distributions of the 2D geometries are derived from the updated pair representation via linear layers and softmax operations. To predict the 1D geometry, we transform the MSA representation into 1D representation by row-wise weighted summation, followed by linear layers and softmax operations to obtain the predicted probabilities.

#### Step 3. Generation of full-atom structure models

Similar to trRosetta, trRosettaRNA generates full-atom structure models by energy minimization with restraints from the predicted geometries.

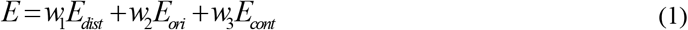

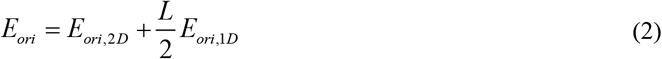

where *E*_*dist*_, *E*_*ori*_, and *E*_*cont*_ represent the distance-, orientation-, and contact-based restraints, respectively; *E*_*ori,2D*_ and *E*_*ori,1D*_ represent the restraints from 2D and 1D orientations, respectively; *L* is the length of the sequence. A detailed description of these energy terms is available in the Supporting Information. The weights (*w*_1_=1.03, *w*_2_=1.0, *w*_3_=1.05) are decided on hundreds of RNAs randomly selected from the training set to minimize the average RMSD. Note that we only select a subset of restraints with probabilities higher than a specified threshold (0.45, 0.65, and 0.6 for distances, orientations, and contacts, respectively).

The folding procedure is implemented with pyRosetta ^36^. From each RNA, 20 full-atom starting structures are first generated using the *RNA_HelixAssmebler* protocol in pyRosetta ^36^. The Quasi-Newton-based optimization L-BFGS is then applied to refine these structures by minimizing the total energy, resulting in 20 refined full-atom structure models. Finally, the model with the lowest total energy (Eq. 1) is selected as the final prediction.

### 2. Construction of datasets

#### Test sets

Two benchmark datasets are constructed in this work. The first one is from the RNA-Puzzles experiments. This set consists of all RNA-Puzzles targets from PZ1 through PZ33 except PZ2. PZ2 is a complex that has complicated interactions among eight chains, which is out of the prediction scope of the current work. The second dataset comes from PDB. In detail, we first collected 339 RNA structures from PDB that are released after 2017-01. RNAs with more than 200 or less than 30 nucleotides were removed. Then the program cd-hit-est ^37^ was used to remove redundant sequences at 80% sequence identity. The final set consists of 101 RNAs.

#### Training sets from PDB

To train our models, we first collected all the RNA chains released before 2022-01 in PDB. Multi-chain structures were separated into single-chain structures. Modified nucleotides are replaced by the standard ones. In addition, if two chains form more than three base-pairing interactions, they are linked by three Adenines, resulting in a new sample. In total, we obtained 8849 samples. Then we tried to generate MSA for each query sequence and removed the sequences without sequence homologs. Finally, 3633 RNA chains were retained for training the network models of trRosettaRNA.

To avoid data leakage in the benchmark tests while keeping as many training samples as possible, five training subsets were obtained by filtering the above 3633 RNA chains. Specifically, for the RNA-Puzzles set, we split the 30 RNAs into four subsets according to their release dates in PDB (i.e., 2010-12~2013-07, 2013-07~2016-07, 2016-07~2019-04, and after 2019-04). Correspondingly, four smaller training sets (1133, 1528, 2337, and 3001 samples, respectively) were obtained by removing structures that were released after the above dates. We trained four network models with these training sets, respectively. For each group of the RNA-Puzzles targets, the predictions were made by the model trained on the corresponding training set. For the 101 test RNAs, the training set consists of 2454 RNAs that are released before 2017-01.

#### Self-distillation training set from bpRNA

As the number of available RNA structures is limited, inspired by the success of the self-distillation method used in AlphaFold2, we constructed a self-distillation dataset from the bpRNA database with experimental secondary structures ^38^. In detail, we collected the bpRNA sequences that are available in the Rfam database ^31^ so that the Rfam MSAs can be used immediately. Then we removed the orphan families (i.e., with one RNA sequence only) and ran cd-hit-est to exclude the redundant sequences at a sequence identity cutoff of 80%. The final self-distillation dataset consists of 13202 RNA chains.

We use a single un-distilled RNAformer model, i.e., trained on the PDB dataset, to generate the predicted labels for the self-distillation set. Using this un-distilled model, we predicted the 1D and 2D geometries (in the form of probability distributions) for every sequence in the self-distillation set. These predicted geometries are then assigned as the labels of these distillation samples. As the predictions may be inaccurate for some nucleotides, we estimate the prediction confidence and filtered out the potentially inaccurate nucleotides and nucleotide pairs. In detail, for each pair of nucleotides (*i, j*) with sequence separation less than 128 (i.e., |*i*-*j*| ≤ 128), we computed the mean P-P distance distribution (i.e., the reference distribution, denoted by 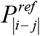), using the predicted distance maps for 1000 samples randomly selected from the self-distillation set. Then for each pair of nucleotides in a self-distillation sequence, we calculated its confidence score (denoted by *c*_*i,j*_), defined as the Kullback-Leibler divergence between its predicted distribution (denoted by *P*_*i,j*_) and the reference distribution:

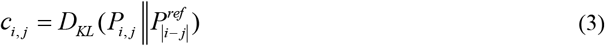

The per-nucleotide confidence score *c*_*i*_ was calculated as the average of *c*_*i,j*_ over all *j*s within the sequence separation of 128:

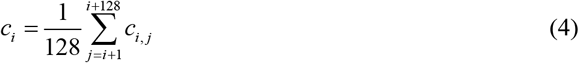

During training (see below), the nucleotides/nucleotide pairs with confidence scores < 0.5 are masked out when calculating the 1D/2D losses, respectively.

### 3. Training procedure and loss function

The training of an RNAformer model can be divided into three steps. In the first step, we trained an un-distilled model using the PDB set by 15 epochs. This model was then used to generate the labels for RNAs in the self-distillation set. In the second step, the un-distilled model was further trained on the combination of the PDB set and the self-distillation set with another 15 epochs. At each epoch, the training samples consist of all the *N* samples from the PDB set and randomly selected 3*N* samples from the self-distillation set, where *N* is the size of the PDB set. In the third step, we finetuned the models on the long sequences (>100 nucleotides) selected from the PDB set. We used the Adam optimizer to minimize the loss function (see below) with different learning rates (0.0001 for the first two steps, 0.00005 for the third step).

For all training steps, the loss function is defined as the cross entropy between the predicted distributions and the real or generated labels. In total, the loss function can be written as:

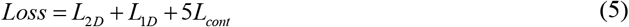

where *L*_2D_, *L*_1D_, and *L*_cont_ are the loss items for the 2D distances and orientations, 1D orientations, and 2D contacts, respectively. More specifically, the three loss items can be written as:

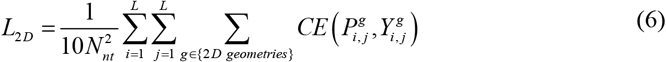

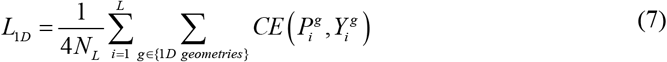

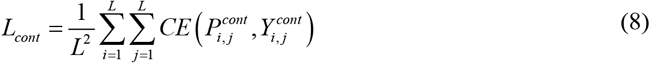

where *CE*() is the cross entropy function; 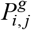 is the predicted probability distribution of the 2D geometry *g* between nucleotides *i* and *j*; 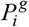 is the predicted probability distribution of the 1D geometry *g* of nucleotide *i*; 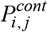 is the predicted probability of nucleotides *i* and *j* to be in contact; the *Y* heads are the one-hot encodings of the true labels (for PDB samples) or the predicted distributions (for self-distillation samples); *L* is the number of nucleotides in sequence; 10 and 4 are the number of types of 2D geometries (5 distances + 5 orientations) and 1D geometries (4 orientations), respectively.

### 4. Estimation of model confidence

To estimate the quality of the predicted model, a few variables are first derived from predicted distance maps and generated decoys.

1) *pRMSD*: the average pair-wise RMSD of the top ten decoys with the lowest total energies.

2) *mp*: the mean probability of the predicted inter-nucleotide distances for the set (denoted by *S*) of the top 15*L* (*L* is the sequence length) nucleotide pairs (as ranked by the probability *P*(*d*_P-P_ < 40Å)). A similar variable has been defined to estimate the accuracy of predicted inter-residue distances ^39^.

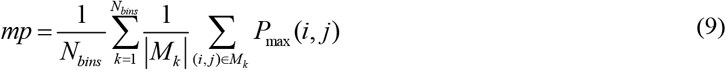

where *d*_P-P_ denotes the distance between the atoms P; *M*_*k*_ is a collection of nucleotide pairs (*i, j*) (from *S*), for which the maximum probability of *d*_P-P_, (i.e., *P*_max_(*i, j*)), belongs to the *k*-th distance bin.

3) *std*, the average standard deviations of the probability values for all nucleotide pairs.

4) *prop*, the proportion of nucleotide pairs with *P*(*d*_P-P_ < 40Å) > 0.45.

The RMSD is estimated based on linear regression over the above variables using hundreds of randomly selected RNAs from the training set.

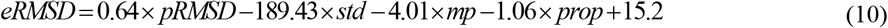

## DATA AND SOFTWARE AVAILABILITY

All datasets used in this work, the trRosettaRNA server, and the standalone package are available at http://yanglab.nankai.edu.cn/trRosettaRNA/.

## ACKNOWLEDGMENTS

This work is supported in part by the National Natural Science Foundation of China (NSFC T2225007, T2222012, 11871290, and 61873185).

