## supporting information for "Accurate *de novo* prediction of RNA 3D structure with transformer network"

**Text S1: Energy function in trRosettaRNA**

trRosettaRNA generates full-atom structure models by minimizing the geometry-derived energy function:

$$E = w_1 E_{dist} + w_2 E_{ori,2D} + \frac{w_2 L}{2} E_{ori,1D} + w_3 E_{cont} \quad \text{(S1)}$$

where $E_{dist}$, $E_{ori,2D}$, $E_{ori,1D}$, and $E_{cont}$ represent the 2D distance-, 2D orientation-, 1D orientation-, and 2D contact-based restraints, respectively; $L$ is the length of the sequence; $w_{1\text{-}3}$ are the weights.

The 2D distances used in trRosettaRNA include the P/C3'/C1'/C/N atom distances, where C/N refers to atom C2/N9 for purine or atom C4/N1 for pyrimidine, respectively. These distances are split into 38 bins, including 37 bins from 3 Å to 40 Å with a 1.0 Å interval and one *no-contact* bin representing the regions <3 Å and >40 Å. trRosettaRNA predicts the probability of each bin and converts the probabilities into the energy potential by the following equation:

$$score^{d}(b) = -ln\frac{P_b + P_B + \varepsilon}{2P_B + \varepsilon} \quad \text{(S2)}$$

where $P_b$ is the probability for the $b$-th distance bin and B is the total number of bins, $\epsilon = 1E\text{-}4$ is the pseudocount parameter to avoid the singularity. Then the distance energy function can be written as :

$$E_{dist} = \sum_d \sum_{i,j \in S_d} score_{i,j}^{d}\left(bin(d_{i,j})\right) \quad \text{(S3)}$$

where $d$ is one of the defined 2D distances; $S_d$ is the set of nucleotide pairs with probability $P$(d<40 Å)>0.45; $d_{i,j}$ is the distance between the $i$-th and $j$-th nucleotides; *bin*() is to convert the distance values into bins.

The 2D orientations used here include two planar angles ($C4'_i$-$N_i$-$C4'_j$, $N_i$-$C4'_j$-$P_{j+1}$) and three dihedral angles ($P_{j+1}$-$C4'_j$-$N_i$-$C4'_i$, $C4'_j$-$N_i$-$C4'_i$-$P_{i+1}$, $N_i$-$C4'_j$-$P_{j+1}$-$C4'_{j+1}$). And the 1D orientations include two planar angles ($P_i$-$C4'_i$-$P_{i+1}$, $C4'_{i-1}$-$P_i$-$C4'_i$) and two dihedral angles ($P_i$-$C4'_i$-$P_{i+1}$-$C4'_{i+1}$ and $C4'_{i-1}$-$P_i$-$C4'_i$-$P_{i+1}$). These planar/dihedral angles are binned into 12/24 segments (15° each) plus one bin referring to the P-P distance <3 Å or >40 Å. The predicted probabilities of angle bins can be converted into the energy potential by the following equation:

$$score^{o}(b) = -ln(P_b + \varepsilon) \quad \text{(S4)}$$

Then the 2D/1D orientation energy functions can be respectively written as:

$$E_{ori,2D} = \sum_{o \in \{2D\ oritentations\}} \sum_{i,j \in S_o} score_{i,j}^{o}\left(bin(o_{i,j})\right) \quad \text{(S5)}$$

$$E_{ori,1D} = \sum_{o \in \{1D\ orientations\}} \sum_{i=1}^{L} score_{i}^{o}\left(bin(o_i)\right) \quad \text{(S6)}$$

where $o$ is one of the defined orientations; $S_o$ is the set of nucleotide pairs with probability $P$(P-distance<40 Å)>0.65; $o_{i,j}$ is the distance between the $i$-th and $j$-th nucleotides; $o_i$ is the 1D orientation corresponding to the $i$-th nucleotide; *bin*() is to convert the orientation values into bins.

The 2D contacts used here were defined as the all-atom minimum distance lower than 8 Å. For the nucleotide pairs with predicted contact probabilities of more than 0.6, we define an additional energy term:

$$cont_{score}{}^{d}(x) = \begin{cases} -5, & x \le d_{cut} \\ -\frac{5}{2}\left[1 - sin\left(\frac{x - \left(\frac{d_{cut}+D}{2}\right)}{D - d_{cut}}\pi\right)\right], & d_{cut} \le x \le 35\ \text{Å} \\ \frac{5}{2}\left[1 + sin\left(\frac{x - \left(\frac{110+D}{2}\right)}{(110-D)}\pi\right)\right], & 35\ \text{Å} \le x \le 110\ \text{Å} \\ 5, & x > 110\ \text{Å} \end{cases} \quad \text{(S7)}$$

where $d$ is one of the distances defined above; $d_{cut}$ is 15 Å for the P-P distance and 20 Å for other distances.

Then the 2D contacts energy functions can be written as:

$$E_{cont} = \sum_{d \in \{2D\ distances\}} \sum_{i,j \in S_c} cont\_score_{i,j}^{d}(d_{i,j}) \quad \text{(S8)}$$

where $d$ is one of the defined 2D distances; $S_c$ is the set of nucleotide pairs with probability $P$(in contact)>0.6; $d_{i,j}$ is the distance between the $i$-th and $j$-th nucleotides.

**Table S1.** Results for 30 RNA-Puzzles targets. The 18 models with RMSD < 4 Å are highlighted in bold.

| RNA-Puzzles ID | PDB ID | Length | $N_{eff}$ | SS Precision | RMSD (Å) | | |
|---|---|---|---|---|---|---|---|
| | | | | | DAS_1 | Best_1 | trRosettaRNA |
| PZ1 | 3MEI | 49 | 16.33 | 1.0 | 3.67 | 3.67 | **2.0** |
| PZ3 | 3OWZ | 84 | 1425.18 | 0.8 | 17.2 | 7.04 | **1.9** |
| PZ4 | 3V7E | 126 | 1551.46 | 0.93 | 4.55 | 3.38 | **2.9** |
| PZ5 | 4P9R | 188 | 5.26 | 0.75 | 10.34 | 10.34 | 14.7 |
| PZ6 | 4GXY | 168 | 7700.42 | 0.78 | 14.33 | 14.33 | 12.9 |
| PZ7 | 4R4V | 185 | 13.13 | 0.94 | 20.63 | 20.63 | 12.4 |
| PZ8 | 4L81 | 96 | 513.52 | 0.83 | 6.2 | 6.20 | **1.8** |
| PZ9 | 5KPY | 71 | 246.04 | 0.83 | 8.89 | 6.27 | 4.7 |
| PZ10 | 4LCK | 96 | 619.87 | 0.7 | 8.42 | 8.42 | **2.2** |
| PZ11 | 5LYS | 57 | 349.25 | 0.85 | 8.63 | 5.87 | 6.7 |
| PZ12 | 4QLM | 125 | 1695.11 | 0.69 | 14.35 | 13.64 | **2.7** |
| PZ13 | 4XW7 | 71 | 409.67 | 0.83 | 7.53 | 7.53 | **2.8** |
| PZ14Bound | 5DDP | 61 | 59.19 | 0.84 | - | 5.79 | **1.8** |
| PZ14Free | 5DDO | 61 | 74.68 | 0.3 | - | 6.98 | 10.3 |
| PZ15 | 5DI4 | 71 | 186.02 | 0.55 | - | 6.98 | 6.1 |
| PZ17 | 5K7C | 62 | 53.66 | 0.78 | 8.83 | 4.9 | **1.4** |
| PZ18 | 5TPY | 71 | 40.09 | 0.88 | 5.36 | 3.16 | **1.3** |
| PZ19 | 5T5A | 65 | 53.94 | 0.94 | 15.31 | 5.54 | 10.0 |
| PZ20 | 5Y87 | 71 | 41.39 | 0.77 | 6.18 | 5.30 | **2.0** |
| PZ21 | 5NWQ | 41 | 20.46 | 0.85 | 5.62 | 3.23 | **1.9** |
| PZ22 | 6JQ5 | 82 | 48.47 | 0.7 | 11.73 | 11.73 | 11.5 |
| PZ23 | 6E8U | 37 | 4.41 | 1.0 | 11.37 | 10.90 | 11.8 |
| PZ24 | 6OL3 | 112 | 24.09 | 0.78 | 16.26 | 10.54 | **3.6** |
| PZ25 | 6P2H | 69 | 677.18 | 0.94 | 5.74 | 2.55 | **3.5** |
| PZ26 | 6PMO | 66 | 1871.27 | 0.94 | 17.21 | 17.21 | **3.0** |
| PZ27 | 6POM | 170 | 2640.27 | 0.89 | 14.87 | 13.27 | 16.3 |
| PZ28 | 6UFM | 98 | 172.62 | 0.8 | 10.89 | 10.89 | **2.5** |
| PZ29 | 6TB7 | 52 | 32.04 | 0.89 | - | 3.51 | **1.6** |
| PZ30 | 7BG9 | 88 | 55.69 | 0.33 | - | 4.43 | 14.6 |
| PZ33 | 7ELP | 46 | 32.42 | 0.84 | 8.22 | 3.64 | **1.4** |
| Average | | 87.9 | 687.77 | 0.8 | - | 7.9 | 5.7 |

**Table S2.** Impact of the different restraints to the structure modeling accuracy on 30 RNA-Puzzles targets.

| **Energy terms** | **RMSD (Å)** |
|---|---|
| 2D distances | 6.1 |
| 2D distances + 2D orientations | 5.9 |
| 2D distances + 2D orientations + 1D orientations | 5.8 |
| 2D distances + 2D orientations + 1D orientations + 2D contacts | 5.7 |

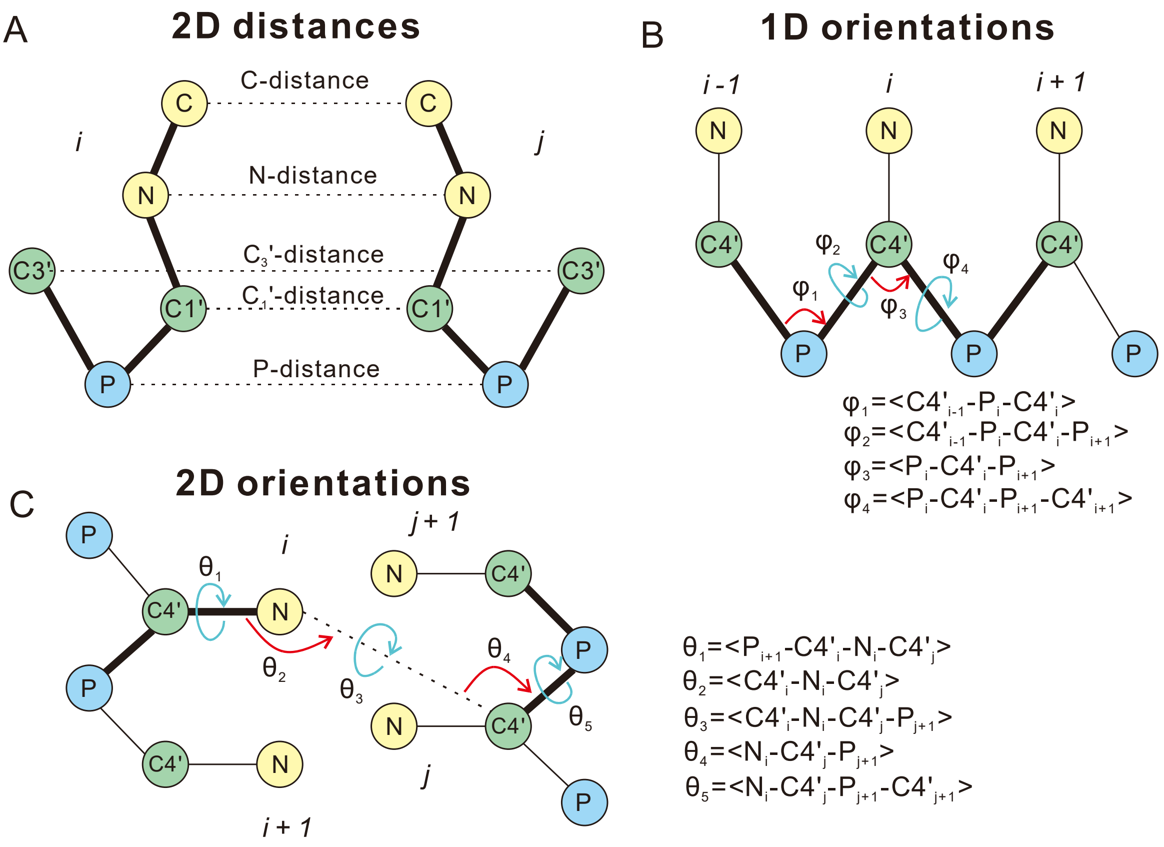


**Figure S1. Definition of the 1D and 2D geometries in trRosettaRNA**. (A) 2D distances. (B) 1D orientations. (C) 2D orientations. N refers to the N9 atom for purine and the N1 atom for pyrimidine. C refers to C2 atom for purine and C4 atom for pyrimidine. *i*, *j* are the indices of nucleotides.

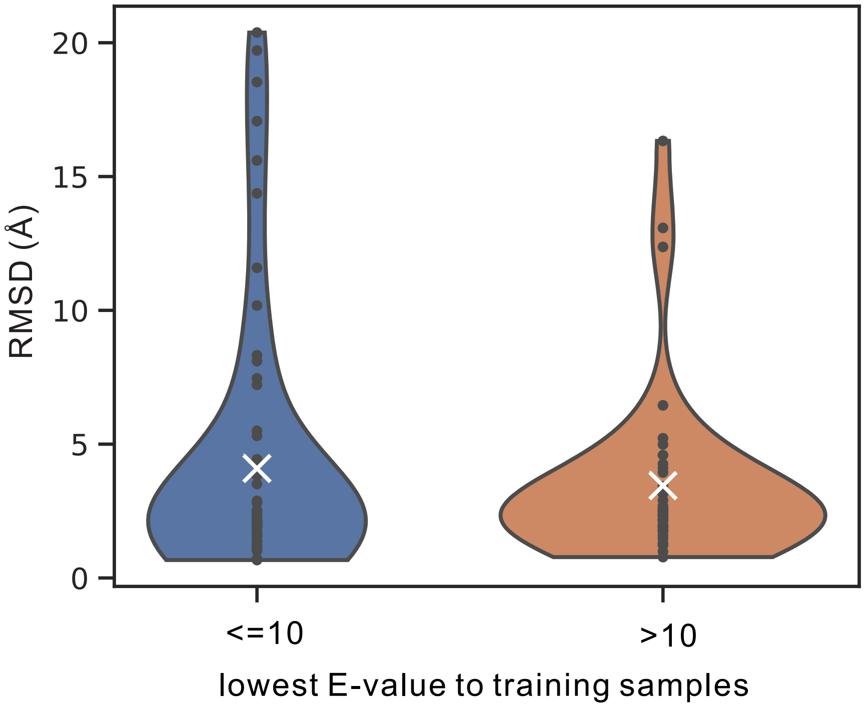


**Figure S2. Relationship between RMSD and the similarity to training test for 101 RNAs.** The white crosses in the violin plots are the mean values.

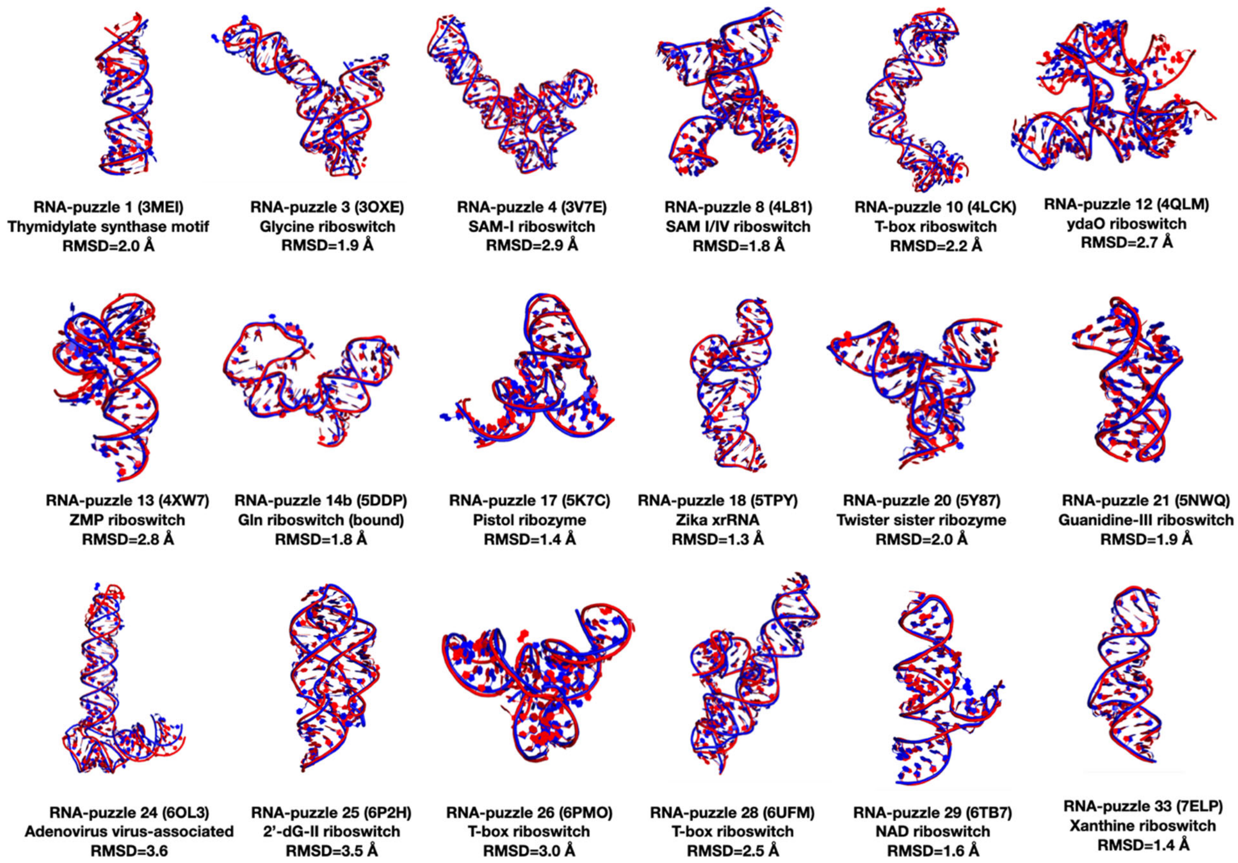


**Figure S3. trRosettaRNA models (red) vs. experimental structures (blue) for 18 RNA-Puzzles targets on which trRosettaRNA achieves <4 Å RMSD.**

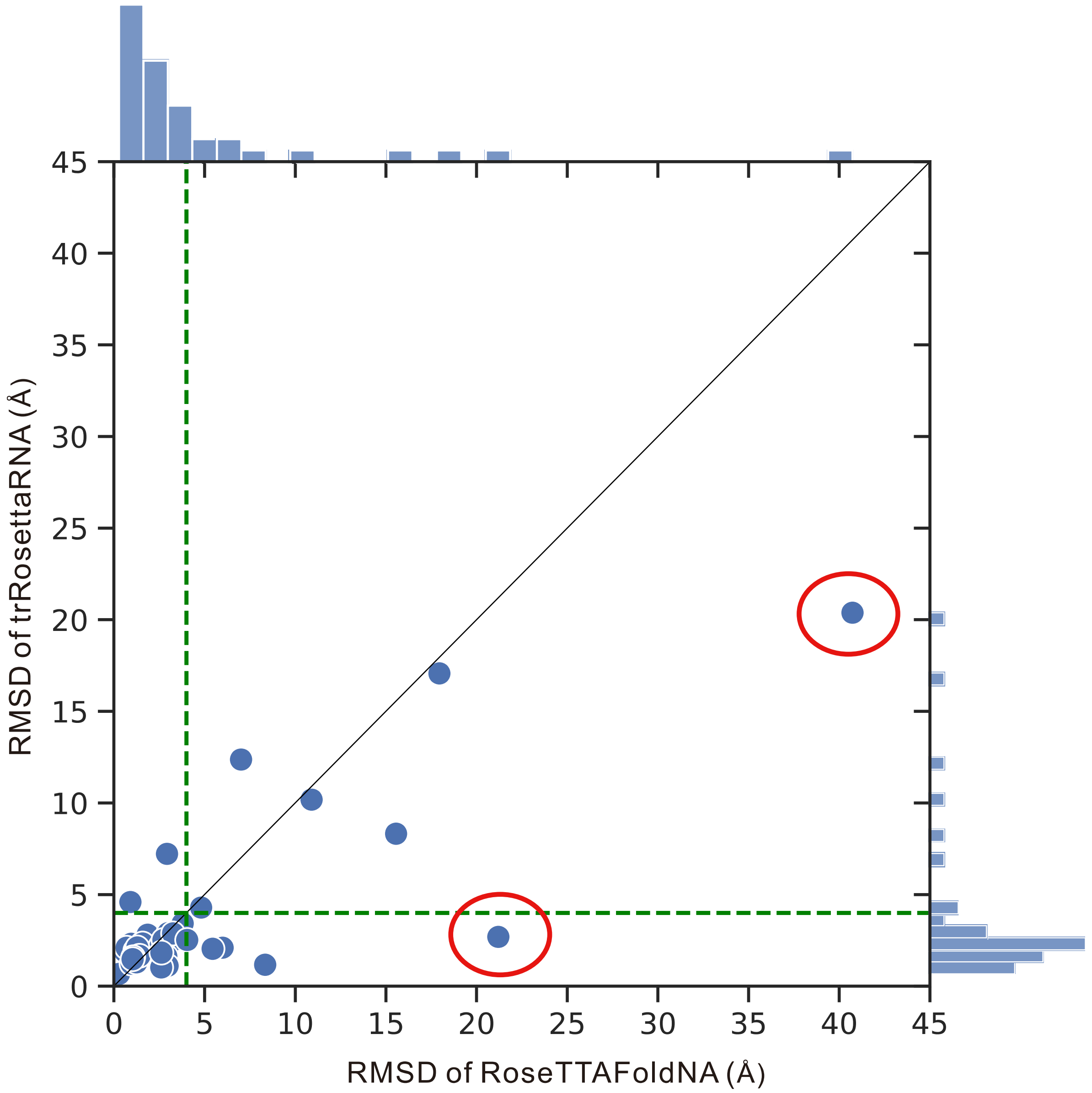


**Figure S4. Head-to-head comparison between trRosettaRNA and RoseTTAFoldNA.** The dashed horizontal and vertical lines correspond to an RMSD of 4 Å. The bar plots show the distributions of RMSDs.

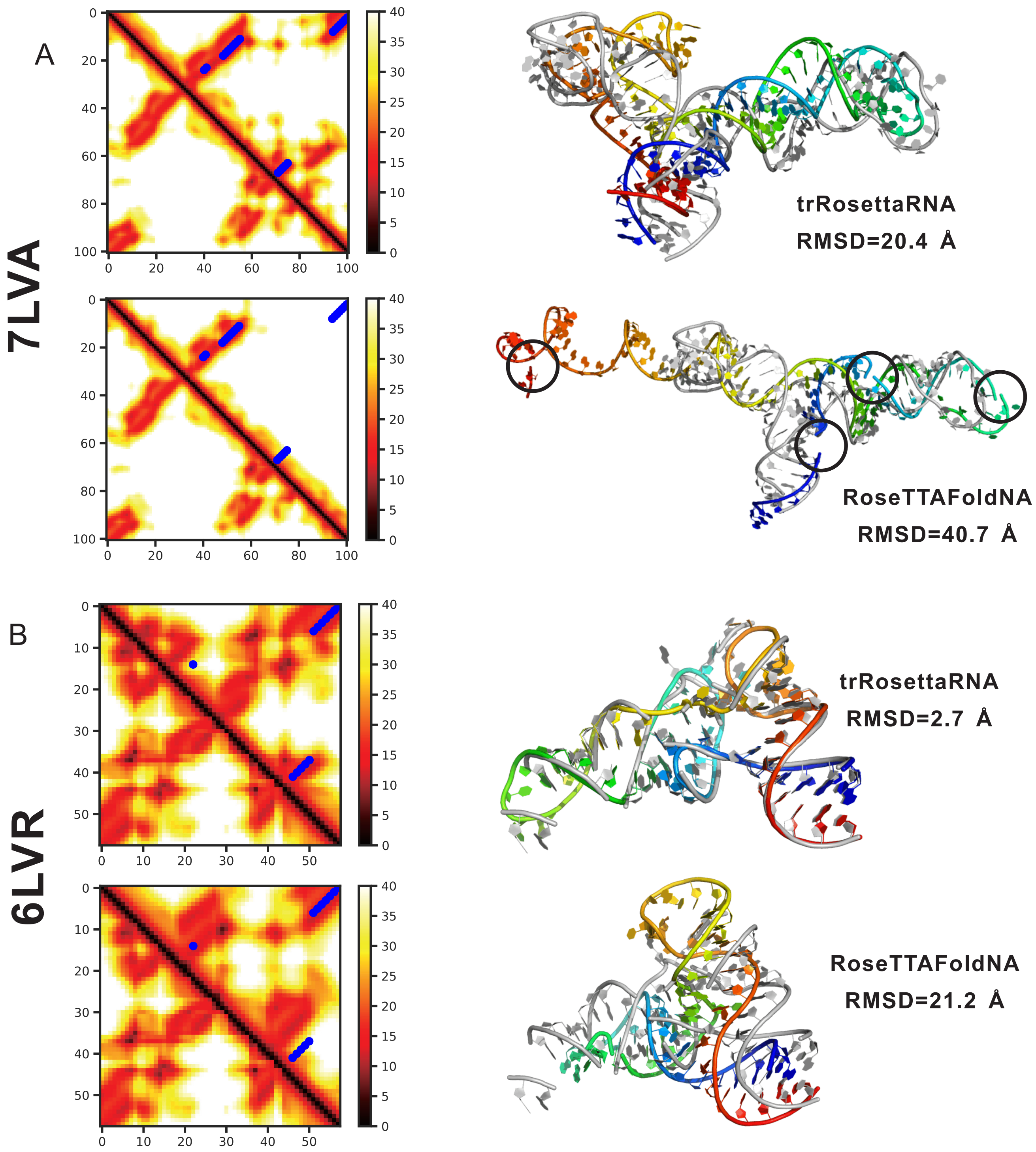


**Figure S5. Comparison between trRosettaRNA and RoseTTAFoldNA on (A) 7LVA and (B) 6LVR.** The upper and lower triangles refer to the predicted and experimental distance maps, respectively. The base pairs predicted by SPOT-RNA are shown as blue points. The predicted structures (rainbow cartoon) are superposed to the experimental structures (gray cartoon).

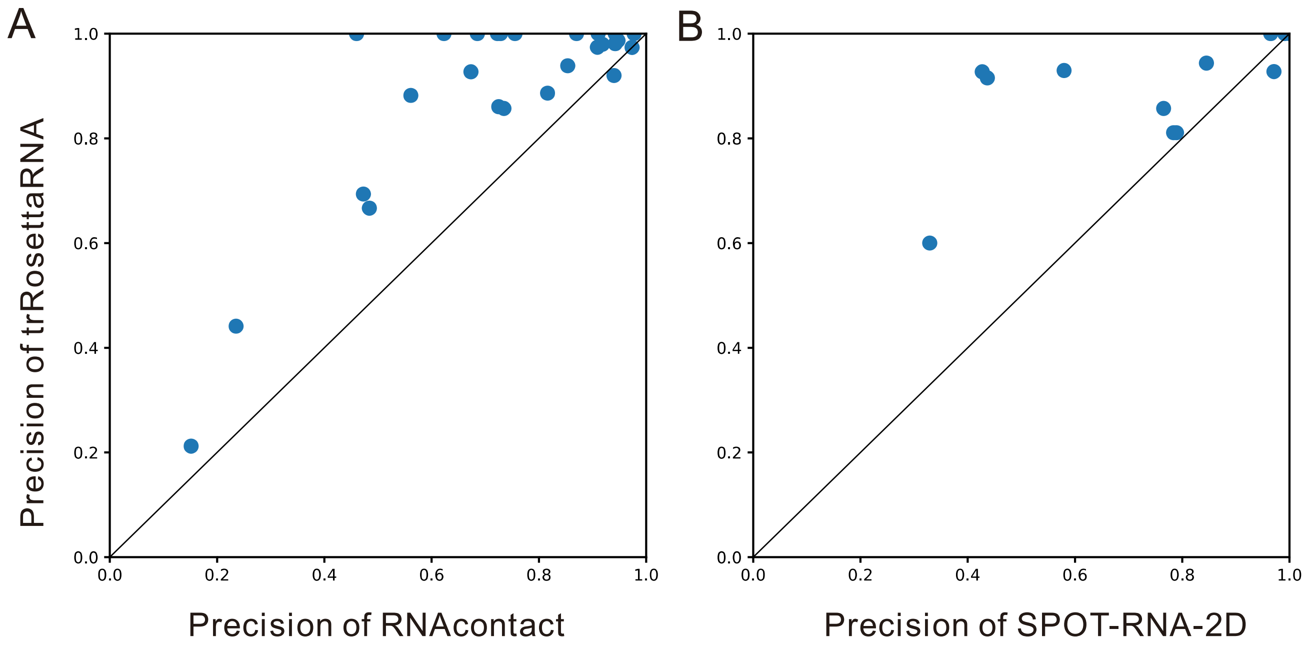


**Figure S6. Comparison of the predicted inter-nucleotide contacts.** (AB) Head-to-head comparison between our method and existing state-of-the-art methods (RNAContact (A) And SPOT-RNA-2D (B)) in terms of the precision of predicted long-range contacts.

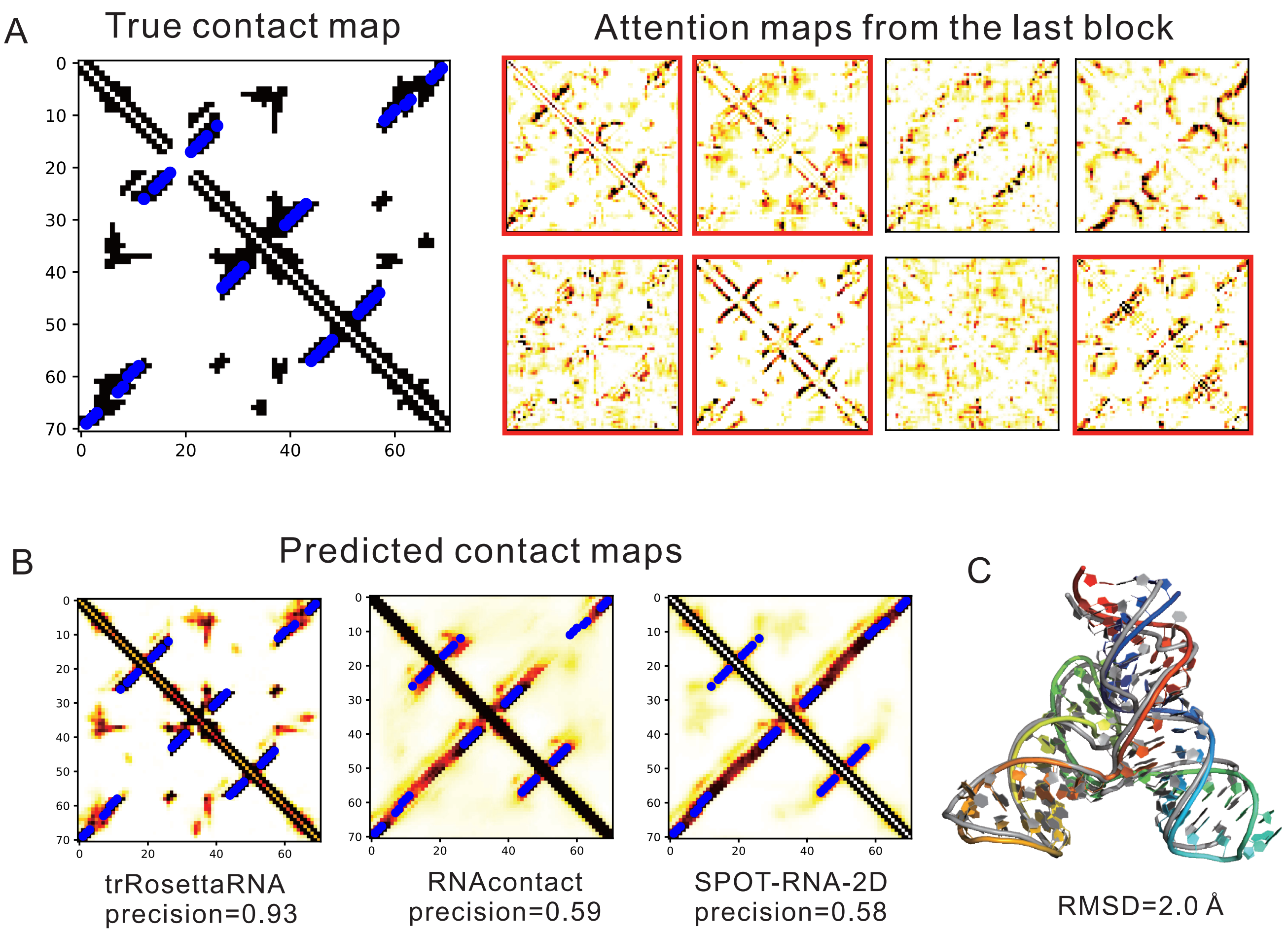


**Figure S7. Analysis of the attention maps produced by RNAformer on the RNA-Puzzles target PZ20.** (A) comparison of the true contact map and the row-wise attention maps extracted from the MSA-to-MSA module in the last block. (B) comparison of contact maps predicted by trRosettaRNA, RNAcontact and SPOT-RNA-2D. (C) the 3D structure predicted by trRosettaRNA. The base pairs are marked as blue points in all contact maps in (A) and (B).

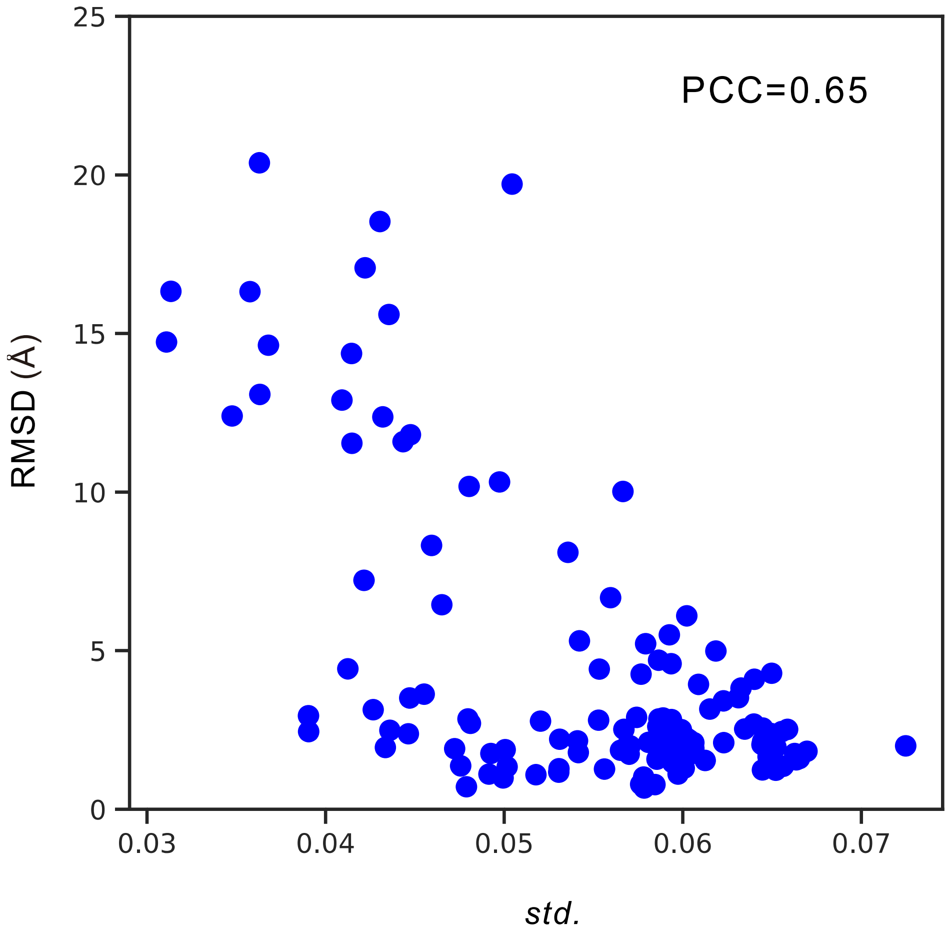


**Figure S8. Relationship between RMSD and the average standard deviations of the predicted distance distributions on the test RNAs.**
